# Age- and estrous-dependent effects of psilocybin in rats

**DOI:** 10.1101/2025.01.10.632408

**Authors:** AL Zylko, RJ Rakoczy, BF Roberts, M Wilson, A Powell, A Page, M Heitkamp, D Fiest, JA Jones, MS McMurray

## Abstract

Psilocybin, a psychedelic compound in “magic” mushrooms, has promise as a novel treatment for psychiatric disorders, many of which are more prevalent in females and have onsets during adolescence. However, there is a lack of research about how factors such as sex and age affect responses to psilocybin, as well as potential safety concerns with developmental exposure. The primary objectives of this preclinical study were to determine if psilocybin-induced head twitch responses differ between adolescent and adult rats, and if estrous phase contributes to variation in female head twitch responses. Secondarily, this study sought to determine if treatment with psilocybin during adolescence has long-term effects on anxiety-associated behaviors and behavioral flexibility. Results showed that 1 mg/kg intraoral psilocybin failed to elicit head twitch responses in adolescents (P35 and P45) but elicited robust responses in adult rats. Further, adolescent psilocybin exposure did not cause long-term differences in performance on the elevated zero maze or probabilistic reversal learning tasks. Lastly, adult females in diestrus showed increased head twitch responses after 1 mg/kg psilocybin compared to females in proestrus. Head twitch responses are thought to be mediated by 5-HT_2A_ receptors, but no age-or estrous-related differences in 5-HT_2A_ receptor expression were observed in the medial prefrontal cortex. Collectively, these results highlight the existence of age-and sex-dependent differences in the effects of psychedelics, while finding no long-term effects on selected behaviors after adolescent exposure. These findings have implications on psychedelic study design, emphasizing the need for inclusive research considering age, sex, and hormonal status.

## 1. Introduction

Psilocybin is one of the primary psychoactive compounds found in “magic” mushrooms (Rakoczy et al., 2024). Consumption of classic psychedelics such as psilocybin can alter consciousness, mood, and cognition. In the past decade, there has been a resurgence of interest in using hallucinogens for the treatment of mood disorders such as major depressive disorder and anxiety disorders. Selective serotonin reuptake inhibitors (SSRIs) are the standard first-line treatment for major depressive disorder and a number of anxiety-related disorders, but require weeks to provide relief and have limited efficacy in many patients (Harmer et al., 2009). This leaves a need for more effective and faster-acting treatments for these conditions. Data from human clinical trials suggests that psilocybin may act faster, require fewer treatments, and have longer-lasting effects than SSRIs (Davis et al., 2021; Raison et al., 2023; von Rotz et al., 2023). Data from preclinical animal studies have largely agreed with these findings and have delved deeper into the potential molecular mechanisms that facilitate these positive outcomes (Lee et al., 2024). However, historically, clinical and preclinical studies assessing treatments for major depressive disorder have been largely relied on adult male subjects (Weinberger et al., 2010), despite the fact that there are higher rates of major depressive episodes in females compared to males (10.5% and 6.2%, respectively), and the past-year prevalence for major depressive episode in adolescents (aged 12-17 years) is 20.1%, which exceeds the reported rates for adults (SAMHSA, 2022). Although there are well-established neurobiological differences between adolescents, adults, males, and females (for example, Koolschijn & Crone, 2013), there is limited research regarding how commonly prescribed medications, such as fluoxetine, may have varying effects across these different populations or lasting influences on behavior after developmental exposure. There are also limited safety and efficacy studies regarding the use of these medications in adolescent and female populations. Identifying potential age- and sex-related differences in psilocybin’s effects is paramount for psilocybin to be effectively used as a therapeutic (Sutherland et al., 2025).

One of the behavioral effects most commonly assessed in preclinical studies of psychedelic drugs, such as psilocybin, is the head twitch response. This behavior, noted across a number of animal species, involves rapid side-to-side and rotational motion of the head or entire body, resembling a wet dog shake (Halberstadt et al., 2020; Halberstadt & Geyer, 2018). When psilocybin is systemically administered to animals, alkaline phosphatase rapidly metabolizes it into psilocin, which crosses the blood brain barrier and activates 5-HT_2A_ receptors in the brain. This activation causes head twitches in a dose dependent manner (Adams et al., 2022; Rakoczy et al., 2024; Thaoboonruang et al., 2024). Evidence for the importance of 5-HT_2A_ activation to produce head twitches stems from studies selectively antagonizing 5-HT_2A_ receptors concurrently with psilocybin administration which attenuates psilocybin’s induction of head twitches in rodents (Canal, 2018; Hesselgrave et al., 2021; Moreno et al., 2013) and hallucinations in humans (Kometer et al., 2013; Vollenweider et al., 1998). Considering that activation of 5-HT_2A_ receptors appears to be necessary for the head twitch response, it is unsurprising that normal variation in receptor expression, due to factors like age or sex, drives differences in the performance of this behavior. For example, the classic psychedelic mimetic 2,5-dimethoxy-4-iodoamphetamine (DOI) has been shown to induce more head twitches in pre-pubertal mice than in adults (Sun et al., 2022). In addition, DOI increases head twitch frequency in female mice, with greater variability when compared to males. (Jaster et al., 2022). Despite these trends, no studies have examined whether psilocybin’s effects depend on the animal’s age or estrous stage.

Justification for the likelihood of age-related variation in psilocybin’s effects stems from prior reports that therapeutic and behavioral effects of other psychedelic and dissociative drugs differ between adolescent and adult subjects. While there is a lack of literature regarding psilocybin’s effects in adolescents, studies have evaluated other dissociative drugs, including ketamine and 3,4-methylenedioxymethamphetamine (MDMA). In humans, use of ketamine anesthesia in pediatric patients lacked the negative emergence reaction (schizophrenia-like symptoms) noted in adults (Hollister & Burn, 1974). In animals, studies across a variety of behavioral contexts and neurophysiological domains suggests adolescents may have fewer or weaker aversive responses to ketamine and MDMA compared to adults (for review see Bates & Trujillo, 2021), and variation in at least MDMA’s toxicity (Chitre et al., 2020; Teixeira-Gomes et al., 2016). Literature on age-related differences in classical psychedelics such as psilocybin is lacking.

In addition to exploring potential changes in psilocybin’s effects during adolescence, it is crucial to determine if long-term effects from acute adolescent exposure occur. The increase in synaptic density from birth to early adolescence, followed by synaptic pruning in adolescence is consistent across species (Watson et al., 2006). In Long Evans rats, there are decreases in dendritic spine density in the medial prefrontal cortex (mPFC) between postnatal days 35 and 90, corresponding to early adolescence and adulthood in humans, respectively (Koss et al., 2014; McCutcheon & Marinelli, 2009). Serotonergic signaling, in particular, plays a large role in guiding neurodevelopment (Miceli et al., 2013), suggesting that pharmacological modulation of this system during development could have long-term consequences in adulthood. Administration of serotonergic drugs, such as the SSRI fluoxetine, to adolescent rats induce a range of behavior changes in adult rats long after treatment has stopped (Iñiguez et al., 2010). Substantial cross-species evidence suggests that a single dose of psilocybin in adults induces a rapid neuroplastic effect, causing wide-spread and long lasting functional and structural reorganization (Brockett & Francis, 2024; Gaddis et al., 2022; Madsen et al., 2021; McCulloch et al., 2022; Olsen et al., 2022; Reinwald et al., 2023; Siegel et al., 2024; Stoliker et al., 2024; Vejmola et al., 2021) and increasing synaptic density (Shao et al., 2021). The combination of psilocybin’s effect on serotonergic signaling and induction of neuroplasticity suggests adolescent exposure could modulate developmentally-appropriate synaptic pruning, leading to long-term changes in brain development and behavior.

Like age-related variation in psilocybin’s effects, it is also essential to determine the nature of sex-differences in response to the drug. Female mice exhibit more head twitch responses than males when given the same dosage of psilocybin (Farinha-Ferreira et al., 2024) or DOI (Jaster et al., 2022), but it is unclear if this depends on estrous stage. Pharmacological activation of estrogen receptors in female rats has been found to increase 5-HT_2A_ mRNA expression and receptor density in the anterior cingulate, anterior frontal, piriform cortex, claustrum, nucleus accumbens, olfactory bulb, and the lateral dorsal raphe nucleus (Sumner & Fink, 1993), and these effects can be blocked by raloxifene, a selective estrogen receptor modulator (Sumner et al., 2007). In female rats, 5-HT_2A_ receptor expression may also vary across different phases of the estrous cycle (Sumner & Fink, 1997). In the estrous cycle, estradiol reaches a peak during the proestrus phase and is at its lowest value during the diestrus phase (Smith et al., 1975). Variation in 5-HT_2A_ receptor expression due to changes in estradiol warrants further study of head twitch responses in females in different stages of their estrous cycle.

The primary objectives of this study are to determine if the rate of psilocybin-induced head twitch responses differ between adolescents and adults, and if variation in female responses is due to the estrous stage of the animal. The secondary objectives are to determine if behavioral differences relate to differences in 5-HT_2A_ receptor expression, and whether treatment with psilocybin during adolescence has long-term effects on anxiety-associated behaviors and measures of behavioral flexibility. We found that adolescents exhibited fewer head twitches than adults, without age-related differences in 5-HT_2A_ receptor expression in the mPFC. Additionally, we found that females in diestrus exhibited more head twitches than those in proestrus, but no differences in mPFC 5-HT_2A_ receptor expression related to estrous stage. Lastly, we found that a single exposure to psilocybin during adolescence had no long-term effects on anxiety-associated behaviors, behavioral flexibility, or acute responses to psilocybin. Collectively, these results suggest that age- and sex-dependent differences in the effects of psychedelics exist, emphasizing the need for inclusive research considering participant age, sex, and hormonal status.

## 2. Methods

### 2.1. Adolescent Study

#### 2.1.1. Overview of Study Design

A general overview of the study timeline, treatment groups, and major outcomes can be found in Figure 1. Briefly, male and female adolescent rats were given either 1 mg/kg psilocybin or vehicle on postnatal day (P) 35 and 45 and head twitch responses were assessed to determine the acute effects of psilocybin. On P90, these animals underwent elevated zero maze testing to evaluate whether adolescent treatment with psilocybin affected adult anxiety-associated behaviors. Beginning on P93, these animals then underwent training and testing for probabilistic spatial reversal learning, to determine if adolescent treatment with psilocybin affected adult behavioral flexibility. Lastly, starting on P125, these animals were again administered 1 mg/kg psilocybin and vehicle (counterbalanced, separated by 1 week) to determine if adolescent treatment with psilocybin affected adult head twitch responses. Lastly, tissue punches from mPFC were collected from separate groups of male and female untreated animals on P35, 45, and during adulthood (P90) to assess 5-HT_2A_ receptor expression levels at each age. All procedures and protocols were conducted in accordance with the National Institutes of Health’s guidelines for the use of animals in research and were approved by Miami University’s Institutional Animal Care and Use Committee.

**Figure 1.**
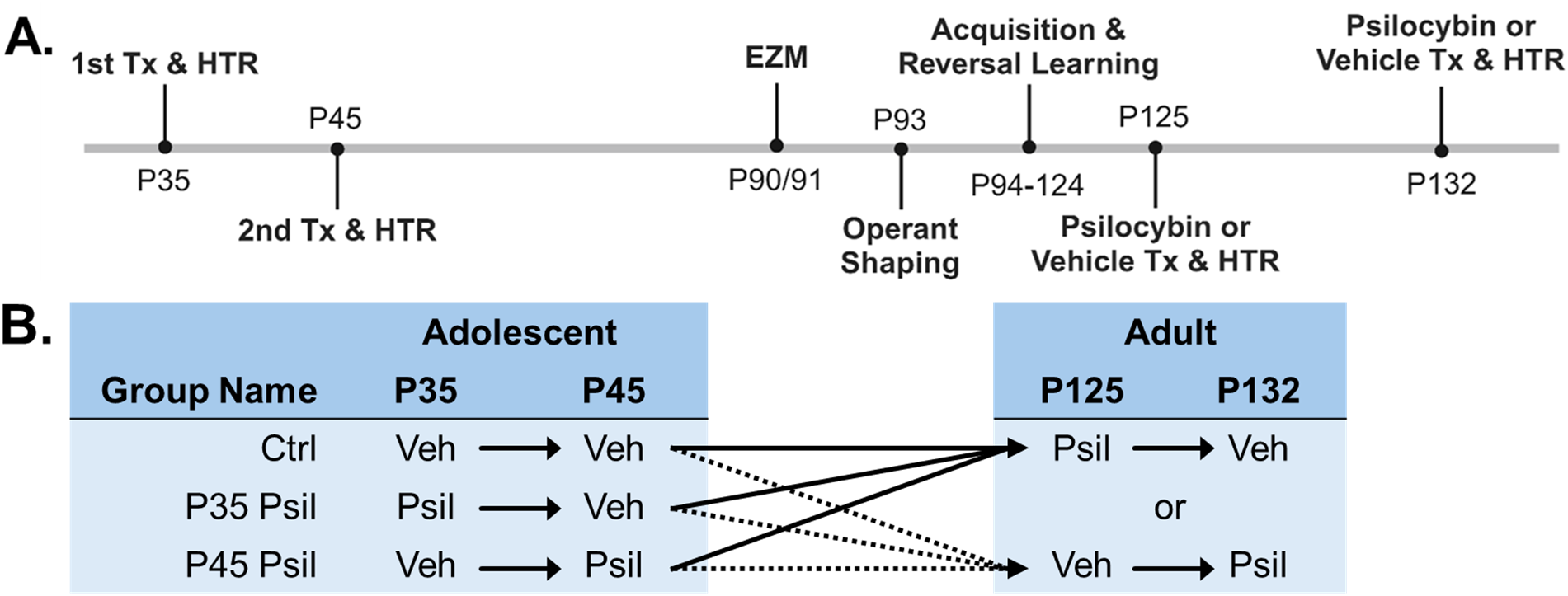
Overview of Adolescent Study. A) Timeline of the study, including treatment (Tx) and behavioral assays (HTR = Head Twitch Response; EZM = Elevated Zero Maze). B) Treatment order and timing for each group.

#### 2.1.2. Subjects

A total of 65 (33 female, 32 male) Long Evans rats were acquired from an in-house breeding colony at Miami University immediately after weaning (P17). Animals received identifying foot tattoos and were pair-housed on P30. They were kept on a 12 hr light/dark cycle (on at 0700 hr) with all experiments occurring during the light cycle (between 0900 and 1400 hr).

#### 2.1.3. Drug Administration and Head Twitch Response Assessment

Psilocybin was synthesized from bioengineered *E. coli* (Adams et al., 2019). Drug was extracted and purified from the cell broth, with purity >98% verified by HPLC and NMR spectroscopy (Adams et al., 2022). Psilocybin (1 mg/ml) or distilled water vehicle was administered by oral gavage of 1 ml/kg. Immediately after gavage, adolescents were placed into a ∼33 cm x 20 cm x 15 cm cage with a transparent lid and video recorded for 30-minutes. Adults underwent identical procedures, except they were placed into a large polycarbonate tube ∼56 cm diameter x 30 cm height for video recording. Head twitch responses in each video recording were then coded by two observers, blind to treatment, with a final concurrence of observers exceeding 90%. Sample video recordings of adolescent and adult head twitch responses can be found in the online Supplement.

#### 2.1.4. Elevated Zero Maze

The elevated zero maze was carried out in a circular testing apparatus (diameter= 40 inches) with two open stretches of maze and two ‘closed arms’ surrounded by 12 inch walls. Tests were run in standard light conditions. Subjects were kept in their home cages for 15 minutes prior to the test in order to allow for habituation to the testing room. After this period, subjects were placed in the center of one of the closed arms and were recorded for 10 minutes. Any-Maze video analysis software (v4.99) was used to calculate total time spent in the open arms as well as total entries into the open arms. The apparatus was cleaned with a 1.5 % quatricide solution between each subject.

#### 2.1.5. Probabilistic Reversal Learning

Methods for the probabilistic reversal learning task have been previously published (Amodeo et al., 2017). Starting one day after completion of the elevated zero maze task (P92), all food was removed from each cage. After this point, subjects were fed once per day with 3.5% of their free-feeding body weight in grams and were weighed daily in order to ensure no drastic weight loss occurred. After twenty-four hours of food restriction, rats began a probabilistic reversal learning paradigm in standard operant chambers (Med Associates, St Albans, VT). Subjects started with a shaping phase where they established that pressing the levers resulted in a sugar pellet reward. Each training session lasted one hour and involved one lever (counterbalanced across subjects) being extended and resulting in a reward. Once the set criterion of at least 30 lever presses was reached two days in a row for each subject, the side of the extended lever switched, and trials continued until this criterion was met again. After a subject was trained on each lever, the acquisition phase began. In this phase, both levers were extended, and subjects learned that one side resulted in a sugar pellet reward 80% of the time, where the other only resulted in a reward 20% of the time. This probabilistic reward aspect was included to make the correct lever slightly harder to establish, consequently making the learning curves more noticeable. The acquisition phase established a “correct” lever for each subject (counterbalanced across subjects); we defined acquisition of the correct lever as the subject choosing the “correct” lever 24 out of 30 possible lever presses two days in a row. After this criterion is met, the side resulting in a sugar pellet reward 80% of the time was switched to the other side. Trials were stopped at 21 days post reversal. The number of days it took the subjects to reach the criterion on the new “correct” lever was recorded, and subjects who never achieved acquisition after reversal were noted as well.

#### 2.1.6. Euthanasia and Tissue Collection

Animals were injected with 0.2-0.5 mL of sodium pentobarbital intraperitoneally and confirmed unconscious via tail and toe pinch before swift decapitation by guillotine. Afterwards, brains were immediately removed and stored whole at -80°C. Brains were then mounted within a cryostat perpendicular to the cutting blade by securing the occipital lobe to a chuck and sectioned until reaching ∼3.5mm anterior-posterior (AP; Paxinos & Watson, 2006). Once the mPFC was identified (0.5 medial-lateral (ML), 3.5 AP, and 3.8 dorsal-ventral (DV)), a 200 µm slice was obtained and a 1mm x 1mm punch of tissue was collected from primarily the prelimbic subregion of the mPFC. Tissue punches were stored at -80°C until use.

#### 2.1.7 5-HT_2A_ Enzyme-linked Immunosorbent Assay

Frozen mPFC tissue punches (∼1 mg of tissue) were rapidly thawed and homogenized using a Dounce homogenizer with tissue suspended in Cell Lysis Buffer (ThermoFisher FNN0011) with a protease and phosphatase inhibitor added (ThermoFisher 78440). Tissue homogenate was then normalized to a concentration of 0.5 mg/mL in lysis buffer using a detergent-compatible Bradford Assay Kit (ThermoFisher 23246) following manufacturer’s protocol. The abundance of the 5-HT_2A_ receptor was then quantified using a commercially available enzyme-linked immunosorbent assay (ELISA) kit (MyBioSource MBS705328). Following manufacturer’s instructions, each sample was tested in duplicate for the quantification of the 5-HT_2A_ receptor. Briefly, 100µL of sample was added in duplicate to wells of a 96-well microplate along with serially diluted standard provided by the manufacturer and incubated at 37°C for two hours. Quantitative analysis was performed by measuring optical density (O.D.) at 450 nm of each well on a microplate reader (BioTek Synergy HT), plotting a standard curve comparing O.D. to known standard concentrations, and then interpolating unknown concentrations for samples using the mean O.D. from duplicate wells. Results are presented as the mean ± standard error of the mean.

### 2.2. Estrous Study

#### 2.2.1. Overview of Study Design

Adult female rats were monitored daily for estrous phase over a two-week period using vaginal lavage to determine baseline estrous cycling and reduce any stress of the lavage procedure. Females then underwent a series of 4 tests (psilocybin and vehicle during both diestrus and proestrus), separated by at least 1 week, in which animals identified as being in diestrus or proestrus were administered psilocybin (1 mg/kg) or vehicle (order randomized) and head twitch responses assessed. Separate groups of untreated animals were euthanized during proestrus and diestrus phases to assess 5-HT_2A_ expression levels in the mPFC. As in the adolescent study, all procedures were approved by Miami University’s Institutional Animal Care and Use Committee.

#### 2.2.2. Subjects

A total of 26 adult (>P100) female Long Evans rats were purchased from Charles River Laboratories (Raleigh, NC). Thirteen females were used for behavioral assessment and 13 were used for assessment of 5-HT_2A_ receptor expression, as described below. All animals were pair-housed and kept on a 12 hr light/dark cycle (on at 0700 hr). Animals underwent all experiments during the light cycle (between 0900 and 1400 hr).

#### 2.2.3. Vaginal Lavage and Estrous Phase Determination

Vaginal lavage samples were collected on clean slides, dried, and stained using the Jorvet Dip Quick Stain Kit (Jorgensen Laboratories, Inc., Loveland, CO) according to the manufacturer protocol. Once dry, slides were examined under a microscope to determine the estrous stage (McLean et al., 2012). Diestrus was characterized by leukocytes with moderate to low cellularity, while proestrus was primarily made up of nucleated epithelial cells.

#### 2.2.4. Drug Administration and Head Twitch Response Assessment

Methods for drug synthesis, administration, and head twitch response testing are identical to those used for adult animals in the Adolescent Study, above.

#### 2.2.5. Euthanasia, Tissue Collection, and ELISA

Methods for euthanasia, tissue collection, and 5-HT_2A_ ELISA are identical to those used in the Adolescent Study, above, except that tissue samples were collected from females during either diestrus or proestrus (instead of specific ages).

### 2.3. Statistical Analysis

Unless stated otherwise, statistical analyses and graphing were accomplished using Graphpad Prism (v8.4). Head twitch responses were analyzed by 2- or 3-way Analysis of Variance (ANOVA), depending on the study and number of conditions. ELISA results and Elevated Zero Maze outcomes in the Adolescent Study were analyzed by 2-way ANOVA. ELISA results for the estrous study were analyzed by unpaired t-test. Prior to analysis, outlier testing was completed on all datasets using Grubbs’ test, and data points identified were removed from the analysis and figures (five instances total across all studies, denoted in results). If significant main effects or interactions were identified in any ANOVA, post hoc t-tests were conducted, with p values corrected for multiple comparisons by the Holm-Sidak method. The rates of acquisition and reversal learning in the Probabilistic Reversal Learning task were analyzed by Cox Proportional Hazards model in SAS (v9.4), with separate survival analyses for acquisition and reversal, and including factors for sex and treatment. Alpha levels for all comparisons were set at p<0.05. Figures include means ± standard error of the mean.

## 3. Results

### 3.1. Adolescent Effects

#### 3.1.1. Head Twitch Response

Acute head twitch responses were quantified for 30 minutes immediately after administration of 1 mg/kg psilocybin or vehicle in early adolescence (P35, n=10-11/treatment/sex), late adolescence (P45, n=10-11/treatment/sex), or adulthood (>P125, n=8-10/treatment/sex). Adolescents receiving psilocybin on either P35 or P45 were compared to age-matched animals receiving vehicle on one (P35) or both days (P35 and P45), respectively. The adult animals included in this analysis also received vehicle treatments on both P35 and P45 (all animals received 4 treatments). Four outliers were identified (1 female and 2 male adult vehicle-treated, and 1 adult male psilocybin-treated animals), and these were removed from further analysis. A three-way ANOVA found significant interactions between age, sex, and treatment [F(2,106)=7.30, p=0.001], age and treatment [F(2,106)=30.26, p<0.0001], sex and treatment [F(1,106)=6.62, p=0.012], and age and sex [F(2,106)=12.51, p<0.0001], as well as main effects of sex [F(1,106)=2.60, p=0.001], treatment [F(1,106)=15.62, p<0.0001], and age [F(2,107)=28.94, p<0.0001]. As shown in Figure 2A, post hoc analyses of comparisons of interest found that compared to vehicle, psilocybin treatment increased head twitch responses in adult females (p<0.0001) and adult males (p=0.0024), but did not increase head twitches in either P35 or P45 animals. Additionally, adult psilocybin-treated females demonstrated more head twitches than either vehicle or psilocybin-treated females on P35 (both p<0.0001) and P45 (both p<0.0001). A similar trend was observed in male animals, with adult psilocybin-treated males exhibiting more head twitches than either treatment group in P35 (both p<0.0005) and P45 males (both p<0.0005). Lastly, adult female psilocybin-treated animals were found to head twitch significantly more than adult male psilocybin-treated animals (p<0.0001).

**Figure 2.**
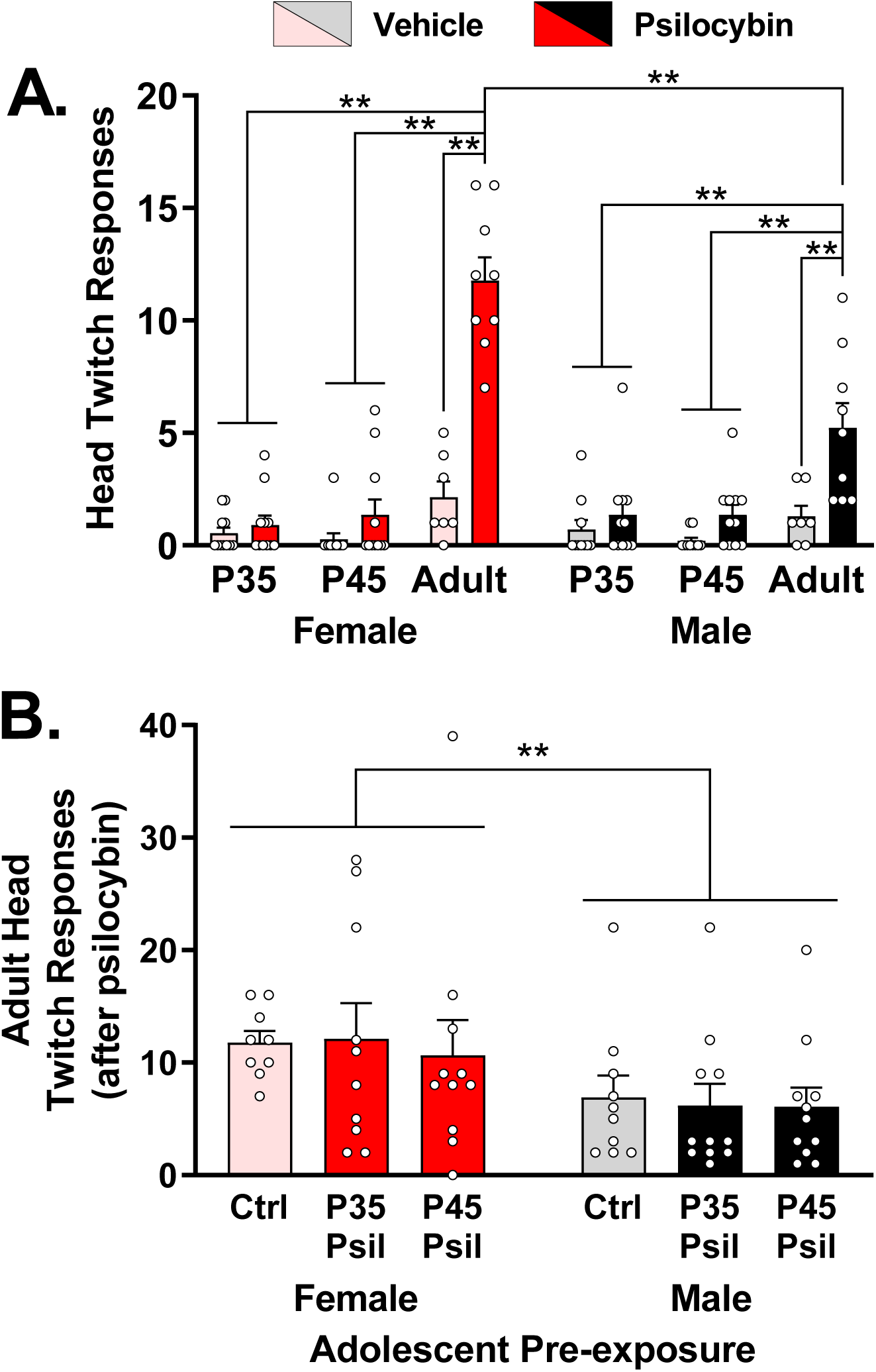
Age-related variation in head twitch responses after 1 mg/kg intraoral psilocybin, and following adolescent pre-exposure to psilocybin. A) Comparison of acute head twitch responses (HTRs) across adolescence and adulthood. Psilocybin failed to increase head twitch responses in adolescents, and adult females exhibited significantly more head twitches than males. Adult subjects depicted received no exposure to psilocybin at any point during adolescence (adolescent Ctrl group). B) Comparison of the effects of adolescent psilocybin exposure on acute HTRs in adulthood. No lasting effect of adolescent pre-exposure to psilocybin was observed on the ability of psilocybin to induce head twitches in adults, a significant main effect of sex was noted with females displaying more HTRs compared to males. Subjects in the Ctrl condition received no psilocybin during adolescence, but received psilocybin in adulthood. Note: **p≤0.01

To determine if adolescent pre-exposure to psilocybin altered psilocybin-induced head twitch responses in adults, animals pre-exposed to psilocybin on P35 or P45 were administered psilocybin again in adulthood (P125 or 132) and head twitch responses assessed. Adult vehicle-treated subjects were not included in these analyses. Two-way ANOVA found no significant effect of adolescent treatment at either time point on adult head twitch responses, but did identify a significant main effect of sex [F(1,56)=7.19, p=0.0096] (Figure 2B).

#### 3.1.2. Adolescent 5-HT_2A_ Receptor Expression

In a separate set of male and female drug naïve animals, tissue samples from the mPFC were collected on P35 (n=2 per sex), P45 (n=2 per sex), or P90 (n=4 per sex) to identify age-related differences in 5-HT_2A_ expression using ELISA. Due to the lack of age-related differences in head twitch responses in adolescence (between P35 and P45), and to reduce the total number of animals used in the study, data was pooled from these age groups. Two-way ANOVA found no significant differences in protein expression due to animal age, sex, or their interaction (Figure 3).

**Figure 3.**
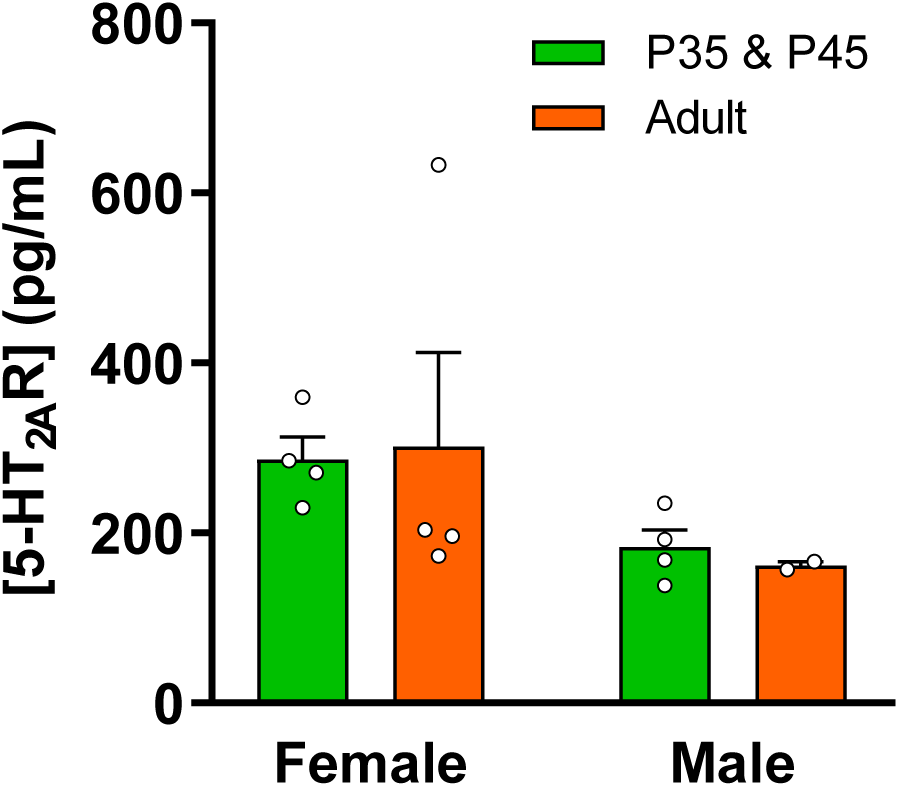
5-HT_2A_ receptor expression in the medial prefrontal cortex of untreated adolescents (P35 & P45) and adults. Quantification of 5-HT_2A_ receptor expression in the medial prefrontal cortex by ELISA demonstrated no significant difference between males and females, adolescents and adults, and no interaction.

#### 3.1.3. Elevated Zero Maze

Animals treated with psilocybin or vehicle on P35 and P45 (n=10-11/treatment/sex/age) completed Elevated Zero Maze testing on P90/91, to determine if adolescent treatment with psilocybin affected levels of anxiety-associated behaviors. Two-way ANOVA of the percentage of time spent in the open areas of the maze (Figure 4A) found no significant effects of sex or treatment, nor their interaction. In addition, no significant effects of sex or treatment were found on the number of entries into the open areas (Figure 4B).

**Figure 4.**
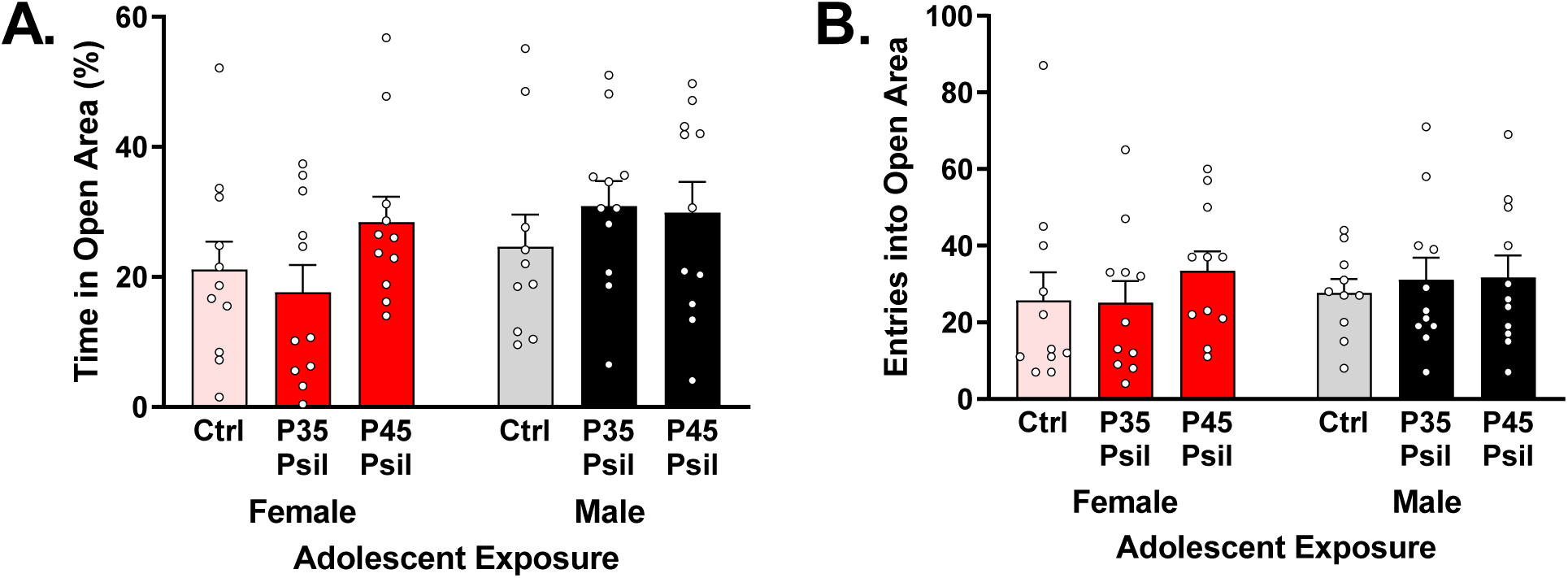
Lasting effects of adolescent psilocybin treatment (P35 or P45) on adult elevated zero maze behaviors (P90/91). A) Percentage of time (600s max) subjects spent in open areas during elevated zero maze test. No significant differences were observed. B) Number of entries into the open areas during the elevated zero maze test. No significant differences were observed. Subjects in the Ctrl condition received no psilocybin at any timepoint.

#### 3.1.3. Probabilistic Reversal Learning

Beginning on P93, animals were trained to complete a probabilistic reversal learning task (30 trials per day), in which one of two possible operant responses (lever press) was rewarded with 80% probability, while the other possible response was rewarded with 20% probability. The rate of initial acquisition of the task (>80% of responses on the higher probability choice) was examined via survival analysis (time to event), which found no significant differences due to subject sex, treatment, nor their interaction (Figure 5A and 5B). Once the basic task was acquired, lever-reward contingencies were switched, and animals continued the task until either >80% responding on the higher probability choice was attained, or 21 days had elapsed. Survival analysis of the rate of reversal learning was conducted, which again found no effect of sex, treatment, or their interaction (Figure 5C and 5D).

**Figure 5.**
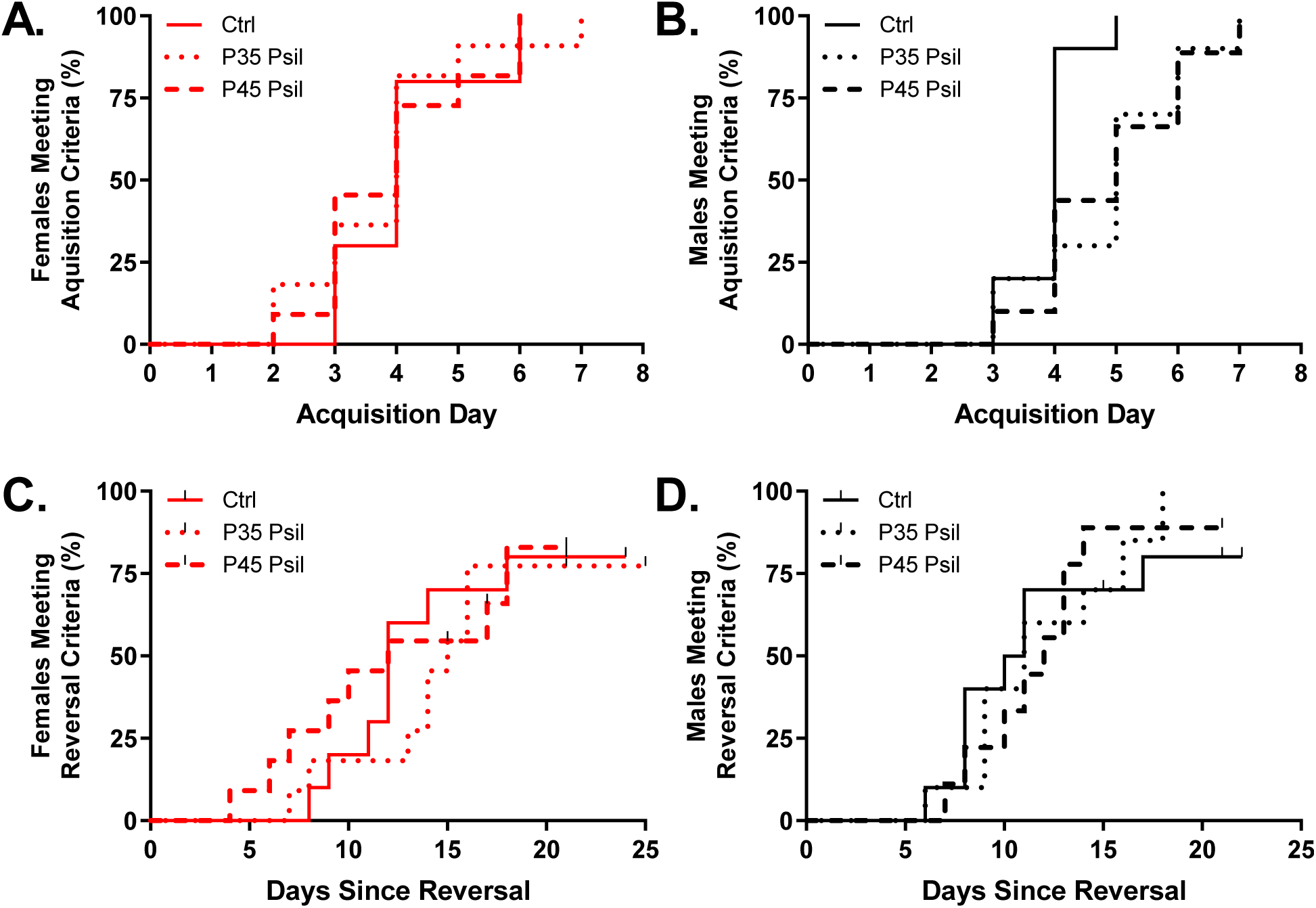
Lasting effects of adolescent psilocybin treatment on adult performance in the probabilistic reversal learning task. The rate of initial task acquisition (criterion of 80% correct responses) did not differ between groups in either females (A) or males (B). There was also no difference in the rate of reversal learning after the operant-reward contingencies were reversed, in either females (C) or males (D). Subjects in the Ctrl condition received no psilocybin at any timepoint.

### 3.2. Estrous Study

#### 3.2.1. Head Twitch Responses

Adult female rats were treated with 1 mg/kg psilocybin or vehicle during the proestrus and diestrus phases of their estrous cycle (4 treatments total per subject), and head twitch responses were quantified for 30 minutes (Figure 6A). One outlier was identified in the proestrus vehicle condition and was removed from the study. Two-way ANOVA found significant main effects of treatment [F(1,45)=32.83,p<0.0001] and estrous stage [F(1,45)=8.396, p=0.0058], but no significant interaction between treatment and stage. Post hoc comparisons found that psilocybin treatment increased head twitch responses compared to vehicle treatment during both proestrus (p=0.0057) and diestrus (p=0.0001), and also that the number of psilocybin-induced head twitch responses is significantly greater in diestrus compared to proestrus (p=0.035).

**Figure 6.**
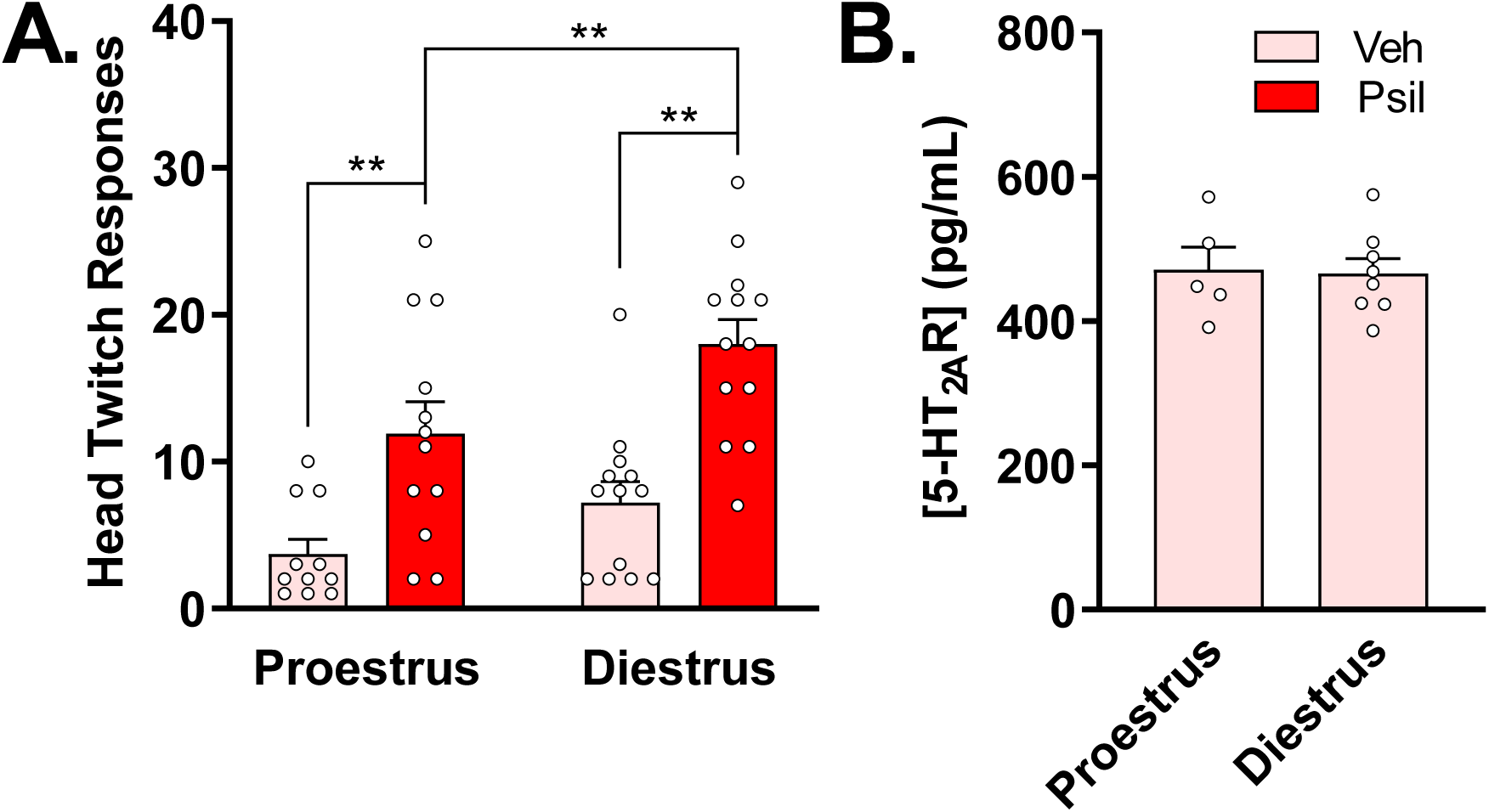
Results from the estrous study. A) Estrous-related variation in head twitch responses in adult females after 1 mg/kg intraoral psilocybin. Females in diestrus exhibited significantly more head twitch responses than females in proestrus. B) 5-HT_2A_ receptor levels in the medial prefrontal cortex in untreated adult females did not differ by estrous stage. Note: **p≤0.01

#### 3.2.2. 5-HT_2A_ Expression

Quantification of the 5-HT_2A_ receptors in the mPFC of proestrus and diestrus adult females was accomplished via ELISA. An unpaired t-test identified no effect of estrous stage on expression [t(11)=0.142, p=0.8900] (Figure 6B).

## 4. Discussion

This study examined the effects of psilocybin on head twitch responses and related behaviors in adolescent and adult, male and female rats, with a goal of identifying potential age-and sex-related variation. Our findings indicate significant differences in psilocybin’s behavioral effects across developmental stages and estrous phases, shedding light on important considerations for therapeutic applications of psilocybin. Psilocybin-induced head twitch responses were pronounced in adult rats, but not in adolescents, with the most pronounced drug effects observed in adult females. Adult females in diestrus exhibited more frequent head twitch responses, suggesting complex interactions between psychedelic effects and the hormonal systems related to estrous.

In adolescents, psilocybin did not induce increased head twitch responses, suggesting a possible developmental difference in drug sensitivity (e.g., different dosage may be needed). The lack of head twitch responses in adolescents aligns with previous studies indicating age-related variation in serotonergic receptor expression and function; however, our data suggest this result was not a consequence of differences in 5-HT_2A_ receptor expression in the mPFC. While age-related differences in 5-HT_2A_ expression in regions not assessed here may be involved, adolescents may also differ from adults in their expression of other serotonin receptor subtypes. Also, methods used for receptor quantification did not distinguish between those in the cellular membrane and those sequestered to intracellular pools, which may differ in their availability to bind with psilocin (Vargas et al., 2023). Furthermore, quantification did not assess the functionality (i.e., ability to activate intracellular signaling cascades) of these receptors. Sequestration of these receptors and their functioning may be partially mediated by effects and concentrations of estradiol *in vivo* (Sumner & Fink, 1995; Compton et al., 2008) and, in part, may explain the lack of congruence between receptor abundance and the behavior observed. Psilocin also has appreciable affinity for 5-HT_2C_ receptors, which have been shown to counterbalance 5-HT_2A_-mediated behavioral effects (Fantegrossi et al., 2010; Van Oekelen et al., 2003), including head twitch responses. It is also possible that other pharmacologic factors, such as age-related variation in the expression of enzymes needed for psilocybin and psilocin metabolism (Saura et al., 1994), could lead to differential metabolism of psilocybin, but this was not the subject of the present study. Recent clinical data has shown that adolescents report similar, and in some cases greater, acute responses to psychedelics (Izmi et al., 2024). Thus, our results do not necessarily suggest that adolescents given psilocybin will fail to hallucinate, but instead suggest the head twitch response may be an unreliable metric for assessing 5-HT_2A_ activation in young rats, highlighting a need for other quantifiable and translatable behavioral measures. Further exploration in other species known to head twitch to a greater degree, and treatment with different dosages than the 1 mg/kg used here, is warranted.

Our study found a lack of significant changes in adult head twitch responses following a single exposure to psilocybin during either early or late adolescence. This suggests that acute exposure during adolescence does not have lasting effects on neural systems responsible for this specific behavior, or that adaptation to those effects has occurred. It is possible that the dosing regimen or the specific developmental time points targeted in this study were insufficient to induce lasting alterations in 5-HT_2A_ receptor expression or function, despite being a behaviorally relevant dose in adults. We also found that a single psilocybin treatment in adolescence did not lead to long-term changes in anxiety-associated behaviors or behavioral flexibility in adulthood. This finding contrasts with reports of other serotonergic drugs, like fluoxetine, which have shown long-lasting effects following chronic adolescent exposure (Cornelius et al., 2005; Iñiguez et al., 2010). The absence of long-term effects may be due to the specific serotonergic mechanisms engaged by psilocybin or the limited duration of exposure used in our study. Regardless, considering that these cognitive domains are thought to underlie the long-lasting therapeutic effects of psilocybin, it is surprising that exposure in adolescence did not cause similar changes.

In adults, the effects of psilocybin on these domains have been shown to persist without supplemental treatment (Carhart-Harris et al., 2016), implying long-term changes in neurophysiology. Thus, the lack of persistent effects in adolescents suggests these drugs may have reduced therapeutic efficacy in this age group, or that the therapeutic effects may have shorter durations. It also indicates that the risk of long-term harm from a single treatment with psilocybin in adolescence may be low. Furthermore, the adolescent brain may be resilient to manipulation of serotonergic signaling, such as produced by psilocybin. Developmental processes leading to synaptic plasticity and pruning during adolescence may overcome any short-term structural changes induced by psychedelics and explain the lack of any chronic effects observed in this study. Adulthood is characterized by reduced synaptic plasticity, potentially allowing for long-term structural changes to persist after psychedelic treatment. For example, induction of synaptic plasticity in brain regions such as the mPFC has been implicated in the action of multiple psychiatric medications (Bittar & Labonté, 2021). Changes in mPFC activity also appreciably alter the activity of downstream afferent targets, which then likely contribute to the hallucinogenic and therapeutic effects of psychedelics. However, the rapid and continual development of these brain regions during adolescence may potentially explain why this early exposure does not cause lasting changes extending into adulthood. Alternatively, these developing afferents may be incapable of generating classical behavioral responses to psilocybin, potentially explaining observed results and more work on this topic is clearly justified.

Alongside the age-related differences and potential long-term effects of psilocybin, the current study also explored whether psilocybin’s effectiveness varied across stages of the estrous cycle. The significant sex differences observed in our adolescent study, particularly the augmented head twitch responses in adult females compared to males, are consistent with the literature indicating greater sensitivity of females to psychedelics (Jaster et al., 2022). Combined with prior reports showing estradiol’s regulation of 5-HT_2A_ receptor expression (Sumner & Fink, 1993), this finding led to our study exploring estrous variation in head twitch responses. Our findings that females in the diestrus phase exhibit more robust head twitch responses compared to females in proestrus provides direct evidence that hormonal fluctuations modulate psilocybin’s effects. Our study was inconclusive regarding the mechanism mediating these effects, however, since we found no differences in 5-HT_2A_ receptor expression across these stages. Although found in similar abundance, it may be that the efficacy of 5-HT_2A_ GPCR signaling depends upon estrous status. Agonism of the 5-HT_2A_ receptor can activate G_q_-mediated signaling pathways, leading to downstream mobilization of intracellular calcium (Kim et al, 2020). Estrous cycle dependent changes to this signaling cascade may alter the ability of psilocin to induce behavioral responses seen normally. Furthermore, 5-HT_2A_ receptor expression was not quantified in other brain regions, and, if variable across the estrous cycle, may mediate the behavioral differences noted here. Future studies should aim to quantify the abundance of the 5-HT_2A_ receptor in other brain regions and further test if intracellular signaling cascades activated by psilocin agonism vary in their functionality across the estrous cycle. Results from this study, at a minimum, underscore the importance of considering the estrous cycle in preclinical studies of psychedelics involving female subjects, and advocate for clinical studies to optimize therapeutic outcomes using gender-specific psychedelic treatment protocols.

The findings of this study contribute to a growing body of evidence that psilocybin’s effects are not uniform across different populations, highlighting the need for more inclusive research that considers age, sex, and hormonal status. Preclinical studies that include both male and female subjects at different stages of development are crucial for generating data that is more translatable to clinical settings. Moreover, understanding the mechanisms underlying age-and sex-related differences in response to psilocybin could inform the development of personalized therapeutic approaches that cater to the unique needs of different patient groups. Our study also raises important questions about the potential long-term effects of psilocybin exposure during critical developmental periods, such as adolescence. Although we did not find evidence of lasting changes following adolescent exposure, future research should explore other behavioral and neurobiological outcomes that may emerge later in life. Future studies should also aim to elucidate the underlying mechanisms driving age- and estrous-related differences and explore their broader implications for psychedelic-assisted therapies, particularly in populations that have been historically underrepresented in clinical trials.

## Supporting information

Video of Adult Male Head Twitch Response

Video of Adolescent Female Head Twitch Response

Video of Adolescent Male Head Twitch Response

Video of Adult Female Head Twitch Response

## Acknowledgements

This work was supported by a sponsored research grant from PsyBio Therapeutics (JAJ and MSM). PsyBio Therapeutics did not contribute to the data collection, analysis, writing, or decision to publish. JAJ is a significant shareholder at PsyBio Therapeutics. JAJ and MSM are co-inventors on patent applications related to tryptamine biosynthesis and use. All other authors declare no conflicts of interest.

